# Cold Receptor TRPM8 as a target for Migraine-associated Pain and Affective Comorbidities

**DOI:** 10.1101/2025.03.25.645275

**Authors:** David Cabañero, Edward P. Carter, Rafael González-Cano, Enrique J. Cobos, Asia Fernández-Carvajal, Antonio Ferrer-Montiel

## Abstract

**Background:** Genetic variations in the *Trpm8* gene that encodes the cold receptor TRPM8 have been linked to protection against polygenic migraine, a disabling condition primarily affecting women. Noteworthy, TRPM8 has been recently found in brain areas related to emotional processing, suggesting an unrecognized role in migraine comorbidities. Here, we use mouse behavioural models to investigate the role of *Trpm8* in migraine-related phenotypes. Subsequently, we test the efficacy of rapamycin, a clinically relevant TRPM8 agonist, in these behavioural traits and in human induced pluripotent stem cell (iPSC)-derived sensory neurons.

**Findings:** We report that *Trpm8* null mice exhibited impulsive and depressive-like behaviours, while also showing frequent pain-like facial expressions detected by an artificial intelligence algorithm. In a nitroglycerin-induced migraine model, *Trpm8* knockout mice of both sexes developed anxiety and mechanical hypersensitivity, whereas wild-type females also displayed depressive-like phenotype and hypernociception. Notably, rapamycin alleviated pain-related behaviour through both TRPM8-dependent and independent mechanisms but lacked antidepressant activity, consistent with a peripheral action. The macrolide ionotropically activated TRPM8 signalling in human sensory neurons, emerging as a new candidate for intervention.

**Significance:** Together, our findings underscore the potential of TRPM8 for migraine relief and its involvement in affective comorbidities, emphasizing the importance of addressing emotional symptoms to improve clinical outcomes for migraine sufferers, especially in females.

## Introduction

Chronic migraine is a painful and debilitating condition that disproportionately affects women. Beyond intense head pain, it is associated with substantial emotional impairments, including anxiety and depression, which are also more prevalent in women and may have a bidirectional relationship with migraine.[1] Additionally, chronic migraine often coexists with confounding conditions like medication overuse headache, which is linked to an altered reward system and heightened impulsivity.[2]

Many migraine triggers have been identified including stress, light or sound stimuli, or menstruation, and while knowledge about pathophysiological mechanisms of migraine has improved in recent years, its pathogenesis remains poorly understood.[1] A few inheritable mendelian forms exist; however, from twin studies, migraine is estimated to have a heritability of 0.4[3], constituting often a polygenic disorder. One of the genes most consistently associated with migraine is *TRPM8.* Thus, single-nucleotide polymorphisms (SNPs) upstream of the *TRPM8* gene are associated with differences in migraine prevalence, particularly in women.[4] This gene encodes Transient Receptor Potential Subfamily M member 8 (TRPM8), a non-selective Ca^2+^ channel receptor initially described in peripheral sensory neurons as sensor of cold, responsive to menthol and its derivatives.[5] Interestingly, TRPM8 has been detected in brain regions kept at euthermic temperature[6] where its physiological role should be different. These include areas such as amygdala and prefrontal cortex which are involved in emotional processing, underscoring potential contributions of TRPM8 to affective dimensions of migraine. This has promoted the present study to explore a role of TRPM8 in common affective traits associated with migraine, using as proxies mouse models of anxiety and depression. In addition, recent identification of TRPM8 activity of classical macrolide compounds like rapamycin and other rapalogs[7, 8], has motivated the experiments presented here to explore new avenues for migraine therapy. These TRPM8 ligands with longer *in vivo* half-lives may offer benefits versus canonical ligands with short-lived effects such as menthol or icilin[9, 10].

Here, we identify emotional behavioural phenotypes in *Trpm8* knockout mice, in naïve conditions and when exposed to a chronic migraine model precipitated by nitroglycerin. We assess rapamycin potential as a modulator of migraine-related behavioural impairments through TRPM8 and validate TRPM8 druggability in human sensory neurons derived from induced pluripotent stem cells (iPSC).

## Results

### An Affective Behavioural Signature of *Trpm8* knockout mice

Humans carrying *TRPM8* SNPs related to migraine susceptibility[4] have such genetic feature during their entire life, hence we speculated that TRPM8-defective mice could serve to identify potential phenotypes altered by distinct channel expression. Thus, we subjected *Trpm8* knockout mice to a battery of tasks to characterise nociceptive and affective behaviour. In agreement with previous literature[5, 8], naïve *Trpm8* null mice showed reduced cold sensitivity (Supplementary Fig.1A). We also observed greater female sensitivity to cold, regardless of genotype (Supplementary Fig.1A). After verification of this reduced cold sensitivity, we aimed to explore affective behavioural traits. First, defensive anxiety was assessed with the Marble Burying task, in which mice are exposed to a number of marbles in a cage with abundant bedding, which mice bury according to their anxious-like phenotype. Wild-type and *Trpm8* knockout mice displayed similar phenotypes (Supplementary Fig.1B). Afterwards, we used the Novelty-Suppressed Feeding test to assess conflict anxiety (Fig.1A). In this test, a hungry mouse is exposed to food in the centre of a novel environment. Mice with higher anxiety levels explore more the surroundings before deciding to bite the pellet in the centre of the arena. Interestingly, male and female *Trpm8* knockouts spent consistently less time exploring the environment and bite the pellet earlier than their wild-type counterparts. This difference could not be attributed to enhanced hunger of *Trpm8* knockouts (Supplementary Fig.1C); hence it could be interpreted either as lower anxiety-like behaviour or as higher impulsivity as previously described.[11] To clarify this, mice were exposed to another classical paradigm of conflict anxiety, the Elevated Plus Maze. In this test, mice choose between exploring the open arms of a plus-shaped maze or staying within the closed arms which offer more safety. Anxious-like animals spend longer time in the closed arms, whereas animals with anxiolytic phenotype are more adventurous and explore more frequently the open arms. In our experimental conditions, we observed similar behaviour among genotypes and sexes (Fig.1A), hence we assumed that differences in Novelty-Suppressed Feeding were due to enhanced impulsivity.

**Fig. 1.**
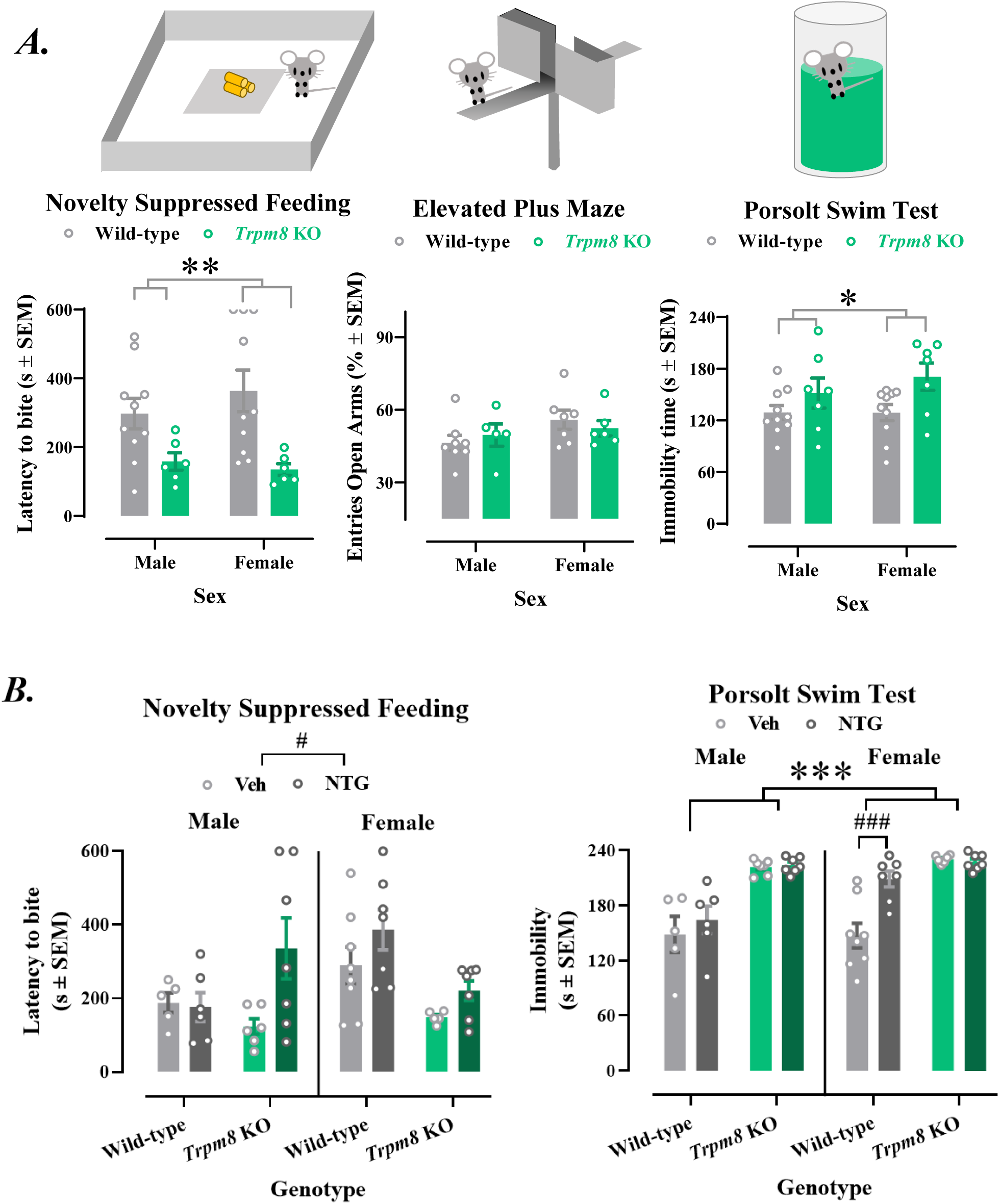
Affective Behavioral Signature of *Trpm8* knockout mice. A) *Affective behaviour of naïve mice.* Left panel, Novelty Suppressed Feeding. *Trpm8* knockout mice of either sex show shorter latency to bite the food pellet in the novel environment than wild-type mice (P<0.05, genotype effect), a phenotype compatible with reduced anxiety-like behaviour or with stronger impulsivity. **Middle panel, Elevated Plus Maze.** Wild-type and *Trpm8* knockout mice of either sex show similar percentage of entries to the open arms of the elevated plus maze, indicating similar anxiety-like behaviour in both genotypes. **Right panel, Porsolt Swim Test.** Male and female *Trpm8* knockouts show longer-lasting immobility than their wild-type counterparts in the Porsolt swim test (P<0.05, genotype effect), indicative of depressive-like phenotype. B) *Affective behaviour of mice subjected to the nitroglycerin model of chronic migraine.* Left panel, Novelty Suppressed Feeding. Mice chronically exposed to nitroglycerin show a significant delay in the latency to bite the food pellet when compared to those exposed to vehicle (P<0.05 treatment effect). This delay is more evident in *Trpm8* knockouts of both sexes and in females, whereas wild-type males appear unaltered. **Right panel, Porsolt Swim Test.** Male and female knockout mice show a robust increase in immobility in the Porsolt swim test, regardless of the pharmacological treatment (P<0.001, genotype effect). In addition, wild-type females exposed to nitroglycerin show a significant increase in immobility (P<0.05 vs. wild-type female vehicle) unlike wild-type males which remain stable, suggesting greater female susceptibility to this depressive-like phenotype induced by nitroglycerin. *P<0.05, **P<0.01, ***P<0.001 Genotype effect, ###P<0.001, #P<0.05 Treatment effect, **A)** 2-way ANOVA; **B)**, 3-way ANOVA; all followed by Tukey. Dots indicate individual values, Error bars are SEM; SEM, Standard Error of the Mean; *Trpm8* KO, *Trpm8* knockout; Veh, vehicle; NTG, nitroglycerin.

After assessment of anxiety-like behaviour, animals were subjected to the Porsolt Swim Test to evaluate depressive-like behaviour. In this test, mice are exposed to forced swimming in an unescapable environment and despair-like behaviour (lack of movement trying to escape) is quantified. Remarkably, this test revealed significant increase of immobility in male and female *Trpm8* knockouts (Fig.1A), suggesting protective effect of *Trpm8* in this depressive trait. In summary, naïve *Trpm8* knockouts displayed reduced cold sensitivity but also enhanced impulsivity and depressive-like behaviour, unaffected by sex.

### Enhanced affective vulnerability of *Trpm8*-deficient mice exposed to the model of chronic migraine

We previously described enhanced mechanical sensitivity of female mice when compared to males after chronic treatment with nitroglycerin, a trigger of migraine-like pain in sensitive individuals.[12] In the same study, we also observed that males became as sensitive as females when they lacked TRPM8[12], revealing a protective function in mechanical hypersensitivity. Here, we replicated these phenotypes (Supplementary Fig.2A) and further evaluated the affective phenotypes altered in naïve *Trpm8* knockouts. Interestingly, in the Novelty-Suppressed Feeding test, mice exposed to nitroglycerin spent longer time before deciding to bite the pellets than mice exposed to vehicle (Fig.1B, P<0.05 Treatment effect), suggesting enhancement of conflict anxiety. This was evident in *Trpm8* knockouts of both sexes and in wild-type females, whereas wild-type males appeared largely unaffected (Fig.1B). Increased latency was unrelated to estimated hunger status (Supplementary Fig.2B). In addition, we quantified depressive-like behaviour with the Porsolt Swim test (Fig.1B). Notably, wild-type females exposed to nitroglycerin exhibited significant depressive-like phenotype (Fig 1B), whereas wild-type males remained unaltered. In contrast, both male and female *Trpm8* knockouts presented prominent despair-like behaviour regardless of vehicle or nitroglycerin exposures. The immobility of vehicle-treated *Trpm8* knockouts was remarkable and enhanced when compared to previous data of naïve *Trpm8* knockouts (Supplementary Fig.2C), suggesting an effect of the experimental paradigm on their depressive-like phenotype. Hence, data indicate protective function of TRPM8, diminishing negative affect and preventing nitroglycerin-induced anxiety and depressive-like behaviour, traits commonly associated with migraine.

### TRPM8-dependent and independent effects of Rapamycin on the murine model of chronic migraine

Following previous reports of rapamycin-induced antinociception in the chronic migraine model[13], wild-type and *Trpm8* knockout female mice received daily low rapamycin doses (1mg/kg) or vehicle during the chronic nitroglycerin treatment (Fig.2A). Rapamycin significantly alleviated nitroglycerin-induced hypersensitivity to mechanical stimulation in wild-type mice, remarkably 2h after the last exposure (Fig.2A, left and AUC). In contrast, this antinociception was absent in *Trpm8* knockouts (Fig.2A, left). Additionally, the delayed long-lasting sensitization after nitroglycerin treatment was reduced in wild-type mice treated with rapamycin when compared to rapamycin-treated *Trpm8* knockouts (Fig.2A, right). This evidenced rapamycin TRPM8-dependent antinociception in the migraine model.

**Fig. 2.**
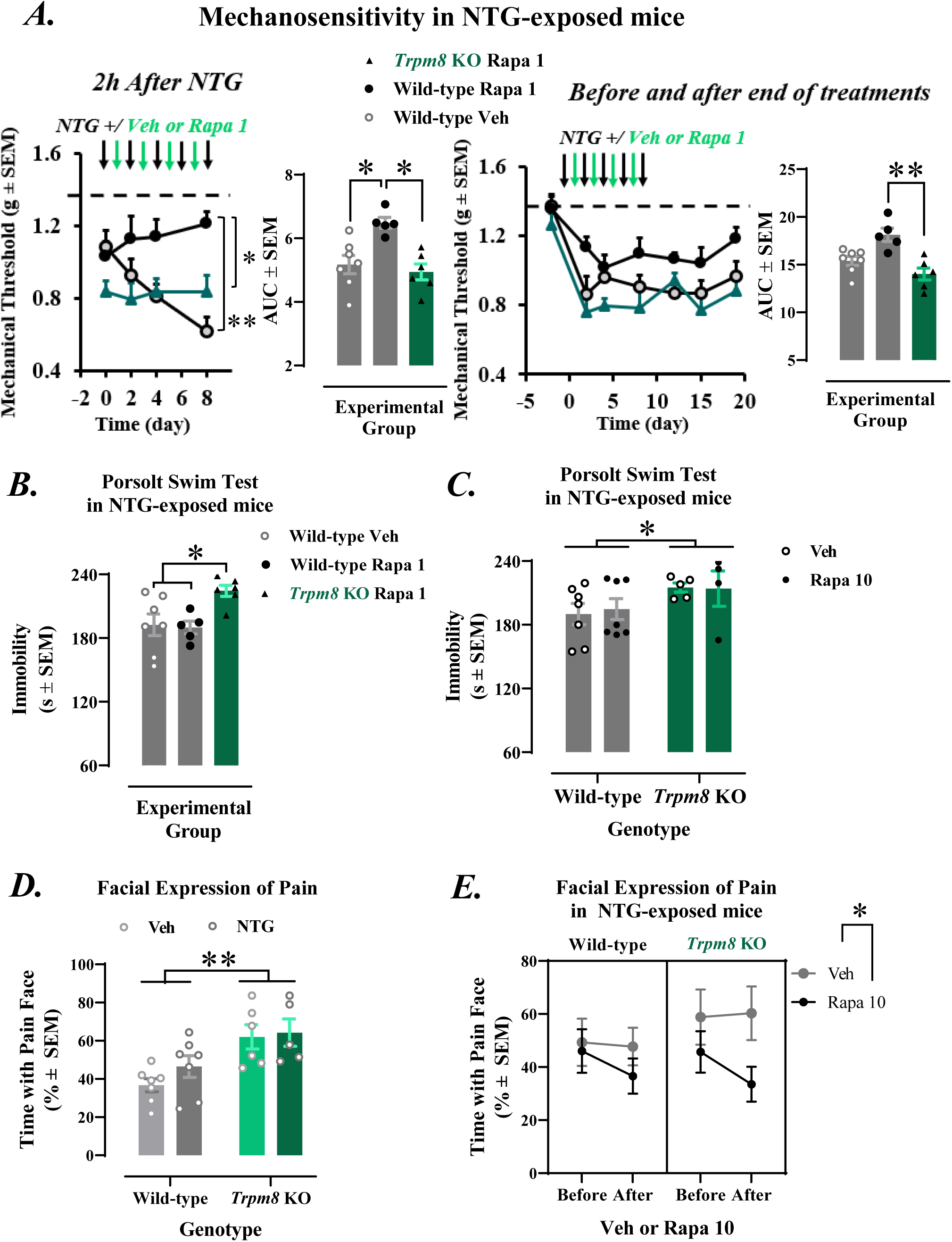
Rapamycin effects on the murine model of chronic migraine. A) Left, Nitroglycerin-exposed wild-type mice treated with low rapamycin doses (1mg/kg, filled circles) show significant alleviation of **mechanical sensitivity** 2h after last nitroglycerin exposure (Left time-course graph, day 8, P<0.05 vs. wild-type receiving rapamycin vehicle, represented by empty circles; left bar graph showing AUCs, P<0.05 vs. vehicle of rapamycin). This rapamycin effect is absent in *Trpm8* knockout mice (vs. green AUCs P<0.05). **Right**, Before each nitroglycerine exposure and after the chronic nitroglycerin treatment, wild-type mice treated with rapamycin show a significant alleviation of mechanical hypersensitivity when compared to *Trpm8* knockout mice receiving the same treatment (right bar graph, rapa wild-type vs. rapa *Trpm8* KO, P<0.01 in AUCs; No significant diference in time course). **B)** After nociceptive measurements in A), mice were subjected to the **Porsolt swim test** after one last additional dose of **1mg/kg rapamycin,** but no effect of this treatment was observed. **C)** Nitroglycerin-exposed mice showed also unchanged depressive-like behavior after repeated treatment with high **rapamycin** doses (**10mg/kg** per day during 4 days). *Trpm8* knockouts preserved their depressive phenotype (P<0.05 vs. wild-type). **D)** Trpm8 knockouts displayed **facial expressions of pain** for longer time than wild-type mice (P<0.05 genotype effect), while nitroglycerin did not enhance this expression of spontaneous pain. **E)** Mice treated with **high rapamycin doses** did show a reduction in the facial expression of pain, regardless of their genotype (P<0.05 Rapamycin treatment effect). NTG, nitroglycerin. KO, knockout. Rapa 1, 1mg/kg rapamycin. Rapa 10, 10mg/kg rapamycin. Veh, Vehicle. SEM, Standard Error of Mean. Error bars are SEM, dots are individual values, bars or dots with error bars are average values. *P<0.05; **P<0.01. Mixed-effects model followed by Tukey post-hoc for time-course. Kruskal-Wallis followed by Dunn’s for 3-group bar graphs, 2-way ANOVA followed by Tukey for 4-group bar graphs. Three-way ANOVA followed by Tukey for rapamycin effect on facial expression.

Once the long-term evaluation of mechanical sensitivity ended, we assessed depressive-like behaviour after one additional injection of 1mg/kg rapamycin (Fig.2B). While depressive-like phenotypes persisted in *Trpm8* knockout mice, rapamycin lacked effect at this low dose (Fig.2B). To explore potential effects of higher doses, we followed a prior study describing rapamycin antidepressant effects, giving 10mg/kg rapamycin once a day 4 consecutive days[14] to nitroglycerin-treated mice. However, depressive-like behaviour of wild-type mice persisted, while *Trpm8* knockout mice kept their enhanced depressive trait (Fig.2C). Hence, this rapamycin treatment was ineffective in reducing depressive-like behaviour associated with chronic migraine, most likely because poor brain distribution.

Finally, we evaluated facial expressions of pain using a computerized neural network[15]. Surprisingly, naïve *Trpm8* knockouts exhibited more facial expressions of pain at rest when compared to wild-type (Supplementary Fig.3). This difference persisted after nitroglycerin (Fig.2D), and nitroglycerin treatment did not modify significantly pain expressions (Fig.2D). Interestingly, nitroglycerin-treated mice exposed to high-dose rapamycin (10mg/kg) significantly reduced the percentage of time with facial expressions of pain regardless of their genotype (Fig.2E). Hence, inhibition of facial expressions of pain was independent of TRPM8. Overall, rapamycin exerted both TRPM8-dependent and independent pain-relieving effects, alleviating nociception and facial expressions indicative of ongoing pain, respectively.

### Activity of Canonical and Novel TRPM8 Ligands in Human Sensory Neurons

Given the protective role of TRPM8 in the murine models, we aimed verification of rapamycin activity in human neurons expressing functional TRPM8. TRPM8 expression and calcium transients elicited after TRPM8 agonists were assessed in human sensory neurons derived from iPSCs. These human neurons showed immunoreactivity to pan-neuronal marker TUJ1 (Tubulin β-III) and BRN3A protein, which peripheral neurons express (Fig.3A). Some of the neurons exhibited also TRPM8 immunolabelling distributed within the cell body (Fig.3B, Supplementary Fig.4A).

**Fig. 3.**
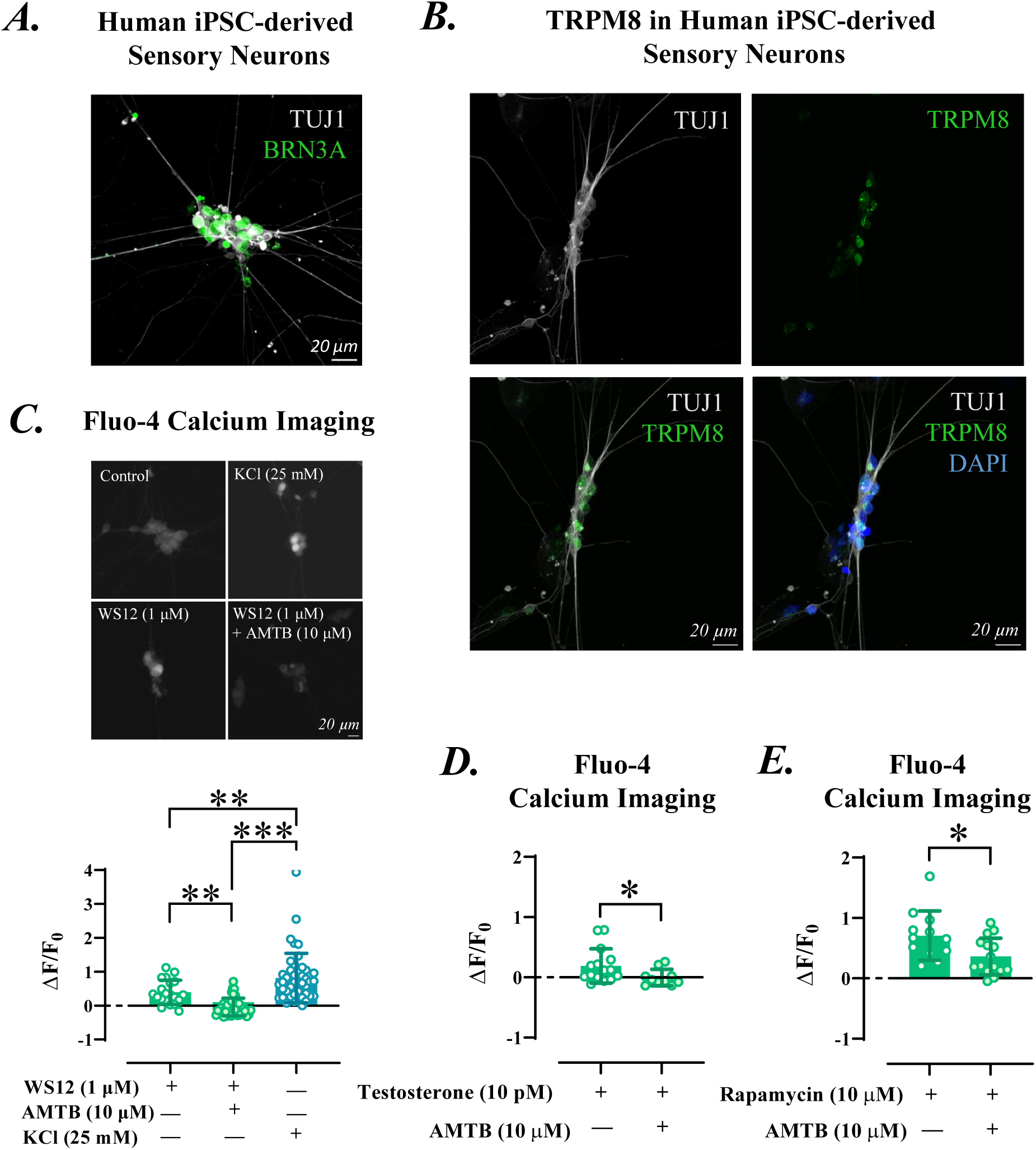
Validation of TRPM8 agonists in human iPSC-derived peripheral sensory neurons. **A)** Human iPSC-derived sensory neurons show immunoreactivity for TUJ1, a β-III tubulin characteristic from neurons (white), and for BRN3A (green), a well-established marker of peripheral neurons. **B)** Part of the human iPSC-derived peripheral neurons also express TRPM8 (green puncta). Nuclei are stained with DAPI (blue). **C)** Fluo-4 calcium imaging conducted in human iPSC-derived sensory neurons show calcium transients in response to 1µM WS12, a selective TRPM8 agonist. These calcium transients are sensitive to the specific TRPM8 antagonist AMTB hydrochloride (P<0.01). **D)** 10 pM testosterone, and **E)** 10 µM rapamycin also elicit AMTB-sensitive calcium transients in human peripheral sensory neurons (P<0.05). Dots indicate individual values and bars are averages normalized to vehicle. Error bars are SEM and scale bars are of 20 µM. *P<0.05, **P<0.01, ***P<0.001. One-way ANOVA followed by Tukey, or Mann-Whitney U.

Functional TRPM8 activity was assessed by intracellular calcium imaging in response to the selective agonist WS12 (1µM), with calcium transients inhibited by the specific antagonist AMTB (10µM) (Fig.3C, Supplementary Fig.4B). Potassium chloride (25mM) elicited pronounced responses, confirming neuronal identity of the cells (Fig.3C, Supplementary Fig.4B). We also tested testosterone (10pM), which evoked modest AMTB-sensitive calcium transients (Fig.3D), consistent with prior findings in murine neurons and HEK cells expressing TRPM8.[12] Finally, rapamycin (10µM[7, 8]) was also evaluated, revealing prominent AMTB-sensitive calcium transients (Fig.3E), which substantiate ionotropic activation of human TRPM8.

## Discussion

The salient contribution of this study is the pronounced influence of the *Trpm8* gene on affective behavioural states relevant to migraine pathophysiology. Namely, TRPM8-defective mice display remarkable depressive-like behaviour, increased impulsivity and frequent facial expressions of pain at rest. When subjected to a chronic migraine model, *Trpm8* knockouts are vulnerable to mechanical nociception[12], and exacerbate anxiety and depressive-like phenotypes. Accordingly, the immunosuppressant and senostatic drug rapamycin, also described as TRPM8 agonist[7, 8], induces TRPM8-dependent relief of mechanical hypersensitivity in mice subjected to the migraine paradigm. Interestingly, rapamycin also inhibits facial expressions of pain through TRPM8-independent mechanisms. Notably, akin to WS12 and testosterone, rapamycin also activates neuronal TRPM8 channels expressed in human sensory neurons, substantiating human TRPM8 engagement in the macrolide activating activity.

The cold hyposensitivity of *Trpm8*-deficient mice[5] is accompanied by a phenotype associated to enhanced impulsivity in the Novelty Suppressed Feeding test.[11, 16] Interestingly, impulsivity was subject of research repeatedly in the context of headache conditions[17] and it involves maladaptations of the reward system linked with medication overuse headache.[2] Such association between TRPM8 and impulsivity could offer mechanistic insight into this poorly understood headache condition.

Naïve *Trpm8* knockouts show depressive-like phenotype in the Porsolt swim test. In agreement, menthol elicited dose-response inhibition in this despair-like behaviour[18] and it can modulate the brain reward system.[19] In line with a disrupted affective phenotype, naïve *Trpm8* knockouts also consistently present more facial expressions of pain at rest than their wild-type counterparts. While full explanation to this unexpected phenotype requires additional experimentation, it is reasonable to expect discomfort in animals with reduced ability to control temperature oscillations.[20] Such a persistent discomfort may be reflected in facial expressions of pain at rest. Overall, TRPM8 appears essential to maintain normal affective phenotypes in naïve mice.

The chronic migraine model promotes behavioural changes in *Trpm8* knockout mice. Males and females exposed to nitroglycerin show prolonged latency to bite the pellet in the novelty suppressed feeding test, indicative of conflict anxiety. Such alteration unveils protective roles of TRPM8 in this emotional trait. In line with this, TRPM8 modulates anxiety-like behaviour at several environmental temperatures[21] and in migraine patients, a *TRPM8* SNP was linked to anxiety prevalence.[22] Interestingly, knockouts exposed to vehicle dramatically enhance despair-like behaviour when compared to naïve knockouts of previous experiments, suggesting a depressive phenotype exacerbated by the experimental paradigm itself. Nitroglycerin-treated wild-type females also develop enhanced depressive-like behaviour, parallel to their mechanical hypersensitivity. Such behavioural repertoire closely mimics the greater vulnerability for depression observed in women with migraine[1]. Indeed, bidirectional influence and significant genetic overlap were observed between migraine and depression.[1] Overall, the data suggests involvement of the *TRPM8* gene in emotional resilience and the interest of exploring TRPM8 modulation for understanding these migraine comorbidities.

Repeated exposure to low rapamycin doses elicits TRPM8-dependent antinociception. The absence of significant effect after the first doses suggests the needs of a minimum concentration to obtain effective antinociception. Since rapamycin half-life is of around 18h in mice[10], it is likely that repeated dosing led to accumulation and significant antinociception by the last day of treatment. Alternatively, this delayed response could be related to TRPM8 signalling downstream of receptor activation.[23] Despite this antinociception, low or high rapamycin doses did not modify depressive-like phenotypes in our experimental conditions. Possible reasons include TRPM8-independent mechanisms, relatively short exposure to rapamycin, or poor blood-brain barrier permeability.[10] Nevertheless, rapamycin antidepressant effects were described in basic studies[14] and clinically in combination with ketamine[24], hence it may be interesting to evaluate rapalogs with higher solubility and brain penetrance, and less secondary effects.[7, 8] The persistence of depressive phenotypes despite alleviation of mechanical hypernociception also suggests independence of the degree of mechanical sensitisation. Hence, sustained hypernociception may be consequence rather than cause of the affective behavioural status. In agreement, other chronic pain models in females showed emotional-like affectation independent of the relief of hypersensitivity[25, 26], and continuous stressful stimuli facilitated mechanical sensitization in females.[27] Overall, our results are compatible with the view that depressive status could be a key factor upstream to the pain sensitization in the behavioural representation of chronic migraine[28]. Rapamycin also inhibits facial expressions of pain in mice exposed to nitroglycerin, independently of TRPM8 signalling. Different mechanisms were proposed for rapamycin-induced antinociception, including mTOR inhibition[29], autophagy modulation[13] or normalization of TNF-α, IL1β and IL-6 levels.[30] In addition, previous reports described improvements in grimace scale face expressions after rapamycin[31], although opposite effects were also observed at high rapamycin doses (20mg/kg[29]). Our data reveal improvement in facial expressions of pain after rapamycin 10mg/kg independently of TRPM8, in agreement with the multiplicity of targets described for this compound.[32]

TRPM8 immunoreactivity and functional expression are revealed in iPSC-derived human sensory neurons. Menthol responses were previously described in these cells,[33] however menthol also evokes TRPM8-unrelated responses.[8] Our data confirms TRPM8-elicited calcium transients after the selective menthol derivative WS12 and sensitivity of those to the specific antagonist AMTB. In agreement with previous works in different cell types[12, 34] we also identify sensitivity of human neurons to testosterone through TRPM8, implying different cold sensitivity between humans of different sex. In addition, human sensory neurons clearly respond to rapamycin and this response is blocked by AMTB. Overall, these results validate previous studies in HEK cells[7, 8] and evidence neuronal responses of human TRPM8 to this naturally-occurring compound.

In summary, we report TRPM8 participation in emotional traits related to impulsivity, anxiety and depression, all of them common migraine comorbidities. In addition, the immunosuppressant and senostatic drug rapamycin is revealed as a primary compound with TRPM8-dependent and TRPM8-independent pain-relieving effects that can modulate the sensitivity of human primary afferent neurons and, therefore, increase the pharmacological armamentarium for chronic migraine intervention.

## Methods

### Animals

Adult male and female mice with a C57BL/6J background (Envigo, Horst, The Netherlands), wild-type or defective in *Trpm8*[5] were bred in the animal facility at Universidad Miguel Hernández (UMH, Elche, Alicante, Spain) and placed in an isolated room in the same institution at least one week before starting the experimental procedures. *Trpm8* knockout mice were a gift from Dr F. Viana (Instituto de Neurociencias de Alicante, Alicante, Spain). Care was taken to minimize the number of animals used and the stress they experienced. Housing conditions were maintained at 21±1°C and 55±15% relative humidity in a controlled light/dark cycle (light on between 8:00a.m. and 8:00p.m.). Animals had free access to food and water except during manipulations and behavioural assessment. Behavioural tests were conducted in a progressive order, starting with the least stressful paradigm and moving to the most stressful, to minimize potential influences between tests when animals were exposed to more than one. The testing sequence began with nociception assessment, followed by the evaluation of anxiety-like behaviour, and concluded with the assessment of depressive-like phenotypes. All procedures were conducted with approval from the UMH Ethical Committee and the regional government (code: 2022VSCPEA0078-2), adhering to European Community guidelines (2010/63/EU).

### Model of Chronic Migraine

Mice received 10 mg/kg nitroglycerin (50mg/50mL Nitroglycerin, Bioindustria LIM, Novi Liguri, Italy) or its vehicle (5% dextrose and 0.105% propylene glycol in pure water) every other day for 9 days (five injections total), administered at 10mL/kg intraperitoneally (i.p.), as previously described.[12]

### Drugs

(1R,2S,5R)−2-Isopropyl-N-(4-methoxyphenyl)−5-methylcyclohexanecarboxamide (WS12; 3040/50, Tocris, Bristol, UK), N-(3-aminopropyl)−2-{[(3-methylphenyl) methyl] oxy}-N-(2-thienylmethyl) benzamide hydrochloride (AMTB; Tocris), testosterone (#T1500, Merck, Darmstadt, Germany) and rapamycin (#J62473.MF, ThermoFisher, Waltham, Massachusetts, United States) were used in cellular studies, all dissolved in 0.01% dimethyl sulfoxide (DMSO, Merck). Used concentrations were based on previous works showing cellular responses in calcium imaging and electrophysiology.[7, 8, 12] In behavioural experiments, rapamycin was administered dissolved in DMSO at 2mL/kg as previously described[35], at a dose of 1mg/kg i.p. for the inhibition of nitroglycerin-induced mechanical sensitisation[13], and at a dose of 10mg/kg to explore inhibition of depressive-like behaviour.[14]

## Behavioural Assessment

### Nociception

#### Cold Plate

Cold response latencies were assessed with a Cold/Hot plate test with a 16.5×16.5cm arena set at 0°C (Bioseb 760112, PanLab, Harvard Bioscience, Cornellà, Barcelona, Spain). Latencies to forepaw withdrawal and licking were recorded as raw latencies in seconds, and cut-off for animal responding was established at 90s.

#### Mechanical Sensitivity

Punctate mechanical sensitivity to von Frey filament stimulation was quantified through the up–down paradigm, as previously reported[12]. Filaments equivalent to 0.04, 0.07, 0.16, 0.4, 0.6, 1 and 2g were used, applying first the 0.4g filament and increasing or decreasing the strength according to the response. Filaments were bent and held for 4–5s against the plantar surface of the hind paws and clear paw withdrawal, shaking or licking were considered nociceptive-like responses. Four additional filaments were applied since the first change of response (from negative to positive or from positive to negative), and the sequence of the last six responses was used to calculate the withdrawal threshold.

### Anxiety-like behaviour

#### Marble Burying Task

Mice were placed in clean translucid cages (42.5×27.5×30cm) with 5cm sawdust bedding overlaid by twenty-eight glass marbles distributed in a 4×7 arrangement. Mice were allowed to explore the cage for 30min and the number of marbles buried (>2/3 of the marble covered by the bedding) was counted.

#### Novelty-Suppressed Feeding

Briefly, animals were food-restricted for 24h and placed in a 51×51cm arena filled with 5cm sawdust, with three food pellets placed in a 12×12cm filter paper situated in the centre. The test ended either when the animal began chewing or when 10min transpired. Immediately afterwards, animals were placed in their home cage and the amount of food consumed in 5min was measured as a relative measure of hunger (mg of pellet consumed).

#### Elevated Plus Maze

Anxiety-like behaviour was evaluated with an elevated plus maze made consisting of four arms (27×6cm), two open and two closed, set in cross from a neutral central square (5×5cm) elevated 40cm above the floor. Light intensity in the open and closed arms was 45 and 5lux, respectively. Mice were placed in the central square facing one of the open arms and tested for 5min. Percentage of entries into the open and closed arms was determined.

#### Depressive-like behaviour

The Porsolt Swim test was used to evaluate depressive-like behaviour.[36] Mice were placed for 6min into transparent Plexiglass beakers (ENDOglassware, 2000mL CBB020, Akralab SL., Alicante, Spain) filled with 1800mL of water at 22±0.2°C to a depth of 22cm. Time of immobility was assessed afterwards for the last 4min. Immobility was considered when the animal made no movements in order to escape (swimming, climbing walls). Water was changed between subjects and beakers cleaned.

#### Facial Expressions of Pain

An artificial intelligence tool, specifically a convolutional neural network trained to analyse video recordings of mouse faces was used to score facial expressions of pain in mice. The procedure has been previously described elsewhere.[15] Briefly, mice were placed individually in custom-made test compartments (50×120×60 mm) with black walls and a mesh-bottomed platform (0.5cm² grid) elevated 1.1 m above the ground. Each compartment was arranged in arrays of four and positioned at the edge of the platform, with one wall open facing a high-resolution infrared video camera (1440×1024 pixels, Kuman RPi camera, USA). Cameras were equipped with two infrared light-emitting diodes and positioned 25cm from the test compartments to encourage the mice to face the visual cliff and, consequently, the camera. Each camera could simultaneously record two mice and was controlled by a Raspberry Pi Zero single-board computer (Kubii, France). Recordings were stored on USB drives for later transfer and analysis. No experimenters were present during testing, except during the first 1–2min when mice were being placed into the compartments. Video recordings lasted a minimum of 15min, but only the 5 to 10min window was analysed to exclude potential artifacts caused by the researcher’s presence at the beginning and to avoid sleep-related features beyond the 10min (e.g., partially closed eyes). The neural network, based on Google’s InceptionV3 model, was trained with facial images of mice highlighting features such as the ears, eyes, cheeks, and nose. We included 245 to clearly exemplify “pain” from animals treated intraperitoneally with cyclophosphamide (300mg/kg), and 300 images labelled as “no pain” from control mice.[37] After over 30,000 training iterations, the model was able to evaluate each frame from new video recordings, assigning a probability value between –1 (no pain) and 1 (pain). We considered that a facial expression denoted pain when the probability value was greater than 0.1. Scripts for DeepLabCut and InceptionV3 were written in Python (v3.5). Network training and video scoring were performed remotely on an Ubuntu Linux computer equipped with an NVIDIA 2080Ti GPU.

### Generation of iPSC sensory neurons

We followed the protocol described previously[33] to obtain sensory neurons from human Pluripotent Stem Cells (hPSCs). Briefly, female human pluripotent stem cells (Healthy Control Human iPSC Line, Female, SCTi003-A, #200-0511, STEMCELL Technologies, Cambridge, UK, CB25 9TL) were cultured under feeder-free conditions using Essential 8 (E8) medium (Thermo Fisher Scientific) on vitronectin-coated plates (Thermo Fisher Scientific). Cells were passaged using 0.5mM EDTA in PBS without calcium or magnesium (Thermo Fisher Scientific) for 5 to 6min to dissociate the hPSC colonies. Cells were maintained in a humidified 5% CO2 atmosphere at 37°C. For neuronal differentiation, hPSCs were plated at 1.5×10⁵cells/cm² on vitronectin-coated 6 well plates in E8 medium containing CEPT cocktail (50nM Chroman 1, 5μM Emricasan, Polyamine supplement (1:1000 dilution) and 0.7μM Trans-ISRIB (#7991, BioTechne)) to improve viability. 24h later cells were switched to Essential 6 (E6) medium (Thermo Fisher Scientific) containing 2µM A83-01 (Transforming Growth Factor-β inhibitor) and 0.2µM CHIR98014 (GSK-3β inhibitor and WNT signalling pathway activator). On day 3, cells were passaged to a single cell suspension with Accutase (STEMCELL Technologies) and seeded at 5.5×10^6^ cells/well in 6 well AggreWell 800 plates (STEMCELL Technologies). Cells were maintained in E6 medium containing 0.5µM CHIR98014, 2µM A83-01, 1µM DBZ (γ-secretase inhibitor), and 25nM PD173074 (FGFR inhibitor). CEPT cocktail was included for the initial 24h during nocisphere formation. On day 14, resulting nocispheres were dissociated using a MACS EB dissociation kit following manufacturers guidelines (#130-096-348, Miltenyi Biotec) and plated on poly-L-lysine/laminin-coated dishes in DMEM/F12 medium, supplemented with N2 supplement, B27 supplement (w/o Vitamin A), 1µM PD0332991 (CDK4/6 inhibitor) and neurotrophic factors (BDNF/GDNF/NGF/NT-3; 25ng/mL each) as described before[33]. By day 28, BRN3A⁺(POU4F1) and Tuj-1⁺ (Tubulin βIII) nociceptor-like neurons were obtained.

### Immunocytochemistry

Cultures were washed with 1X PBS (D8662, Merck) three times. Afterwards, cells were fixed with 4% paraformaldehyde (#28909, Thermo Fisher Scientific) for 20min at room temperature. Permeabilization was achieved with 0.1% v/v Triton 100X (P8787, Merck) for 5min and blocking with 5% bovine serum albumin (#A7906, Merck) for 30min, both in 1X PBS. Cells were labelled with primary antibodies mouse anti-BRN3A 1:100 (#MAB1585, Millipore Sigma, Merck KGaA, Darmstadt, Germany), rabbit anti-TUJ1 1:400 (#5568, tubulin beta-3, Cell Signalling Technology, London, UK) and /or mouse anti-TRPM8 Clone OTI7A11 1:100 (#TA811228S, Origene Technologies GMBH, Herford, Germany) and incubated for 1h at room temperature. Secondary antibodies Donkey anti-mouse-488 1:200 (#A-21202, Thermo Fisher Scientific) and Donkey anti-Rabbit-555 1:200 (#A-31572, Thermo Fisher Scientific) were incubated for 1h at room temperature protected from light. Slides where mounted with ProLong Gold antifade reagent with 4′,6-diamidino-2-phenylindole (DAPI, #P36931, Thermo Fisher Scientific) and images acquired with a confocal microscope (LSM 880, ZEISS, Jena, Germany).

### Calcium Imaging

Fluo4-AM (F14201, Molecular Probes) was dissolved in DMSO at a concentration of 10mM and used at 2µM to load the cells for calcium imaging experiments. D28 iPSC sensory neurons were incubated with Fluo4-AM for 30min at 37°C in standard extracellular solution (in mM: 140 NaCl, 3 KCl, 2.4 CaCl2, 1.3 MgCl2, 10 HEPES, and 5 glucose, adjusted to pH 7.4 with 1M NaOH). Cells were washed three times with extracellular solution and equilibrated for 30 min prior to imaging. Fluorescence measurements were obtained on an LSM 880 confocal fluorescent microscope using a 20x objective. Basal fluorescence was captured every 3s over a 60s period prior to application of stimulants with images captured for a further 3min. Mean fluorescent intensity values were recorded per cell and normalised to pre-simulation. Data expressed as ΔF/F0 using the equation: ΔF/F0=(F_max_-F_0_)/F_0_.

### Statistical Analyses

Behavioural and Calcium Imaging data were analysed using GraphPad Prism9 (GraphPad Software Inc., USA). Sample Sizes were based in previous works evaluating similar behavioural paradigm or calcium transients[12, 16, 26, 35]. For the behavioural experiments in naïve mice, 2-way Analysis of Variance (ANOVA) was used, with factors “genotype”, “sex” and their interaction, followed by post-hoc Tukey recommended tests to compare between experimental groups. When mice were exposed to nitroglycerin, data were assessed with 3-way ANOVA, adding the factor “treatment” and the interactions. Time-courses were analysed with 3-way (Time, Treatment, Genotype) or 2-way (Group, Time) Repeated measures ANOVA followed by Tukey. Calcium Imaging data were analysed with One-way ANOVA (WS12 vs WS12+AMTB vs. KCl) followed by Tukey or with Kruskal Wallis tests (Testosterone vs. Testosterone+AMTB; Rapamycin vs. Rapamycin+AMTB) followed by Dunn’s Multiple Comparison Tests. Outlier. Differences were considered statistically significant when P<0.05. Outliers (±2SD from the mean) were excluded. Artwork was designed using GraphPad Prism, Excel and Power Point. Experimenters were blinded to the key factors being assessed in each behavioural paradigm. Specifically, these factors included genotype in experiments in naïve mice, nitroglycerin or vehicle treatment in the migraine model experiments, and rapamycin vs. vehicle in experiments assessing the effects of rapamycin. In experiments with three experimental groups, only the treatment condition for wild-type mice was blinded. Blinded treatments were allocated randomly through the Random tool of Excel. Raw data and statistical analyses are provided in a Supplementary File (Supplementary_Data_Trpm8.xls).

## Declarations

### Ethics Approval

All procedures were conducted with approval from the UMH Ethical Committee and the regional government (code: 2022VSCPEA0078-2), adhering to European Community guidelines (2010/63/EU)

### Consent for publication

Not applicable

### Availability of data and materials

Raw data and Statistical Analyses corresponding to the Figures shown in this article are provided in a Supplementary File (Supplementary_Data_Trpm8.xls). Videos of the analysed behaviours can be made available under reasonable request.

### Authors’ contributions

D.C. conducted behavioural assays, analysed the data, conceptualized and designed the study and experiments, coordinated *in vivo* and *in vitro* experiments and wrote and edited the first and subsequent drafts of the manuscript. E.P.C. performed cell culture and differentiation assays, executed and analysed calcium imaging and immunocytochemical assays, edited and revised manuscript. R.G.C. developed the neural network and recording device, trained neural network, assisted and trained D.C. in face data management, edited and revised manuscript. E.J.C. developed the neural network and recording device, trained neural network, assisted and trained D.C. in face data management, edited and revised manuscript. A.F.C. supervised and designed experiments, conceptualized the project, revised the manuscript and provided funding. A.F.M. supervised and designed experiments, conceptualized the project, wrote, revised and edited the manuscript and provided funding.

## Supporting information

Supplementary_Data_TRPM8

## List of Abbreviations

AMTB: N-(3-aminopropyl)−2-{[(3-methylphenyl) methyl] oxy}-N-(2-thienylmethyl) benzamide hydrochloride
ANOVA: Analysis of Variance
CEPT cocktail: Chroman, Emricasan, Polyamine and Trans-ISRIB cocktail.
DAPI: 4′,6-diamidino-2-phenylindole
DMSO: dimethyl sulfoxide
E6 medium: Essential 6 medium
E8 medium: Essential 8 medium
HEK cell: Human Embrionic Kidney cell
hPSCs: human Pluripotent Stem Cells
iPSC: induced Pluripotent Stem Cells
SNPs: single-nucleotide polymorphisms
TRPM8: Transient Receptor Potential Melastatin 8
*TRPM8*: Transient Receptor Potential Melastatin 8 human gene
*Trpm8*: Transient Receptor Potential Melastatin 8 mouse gene
TUJ1: Tubulin β-III
WS12: (1R,2S,5R)−2-Isopropyl-N-(4-methoxyphenyl)−5-methylcyclohexanecarboxamide

## Acknowledgements

Authors acknowledge excellent technical assistance of José Manuel Serrano García, Tania Trujillo Ruiz and help of undergraduate students Eva M. Amorós Rojas and Mónica Gamo Muñoz.

## Supplementary Figure Legends

**Supplementary Fig. 1.**
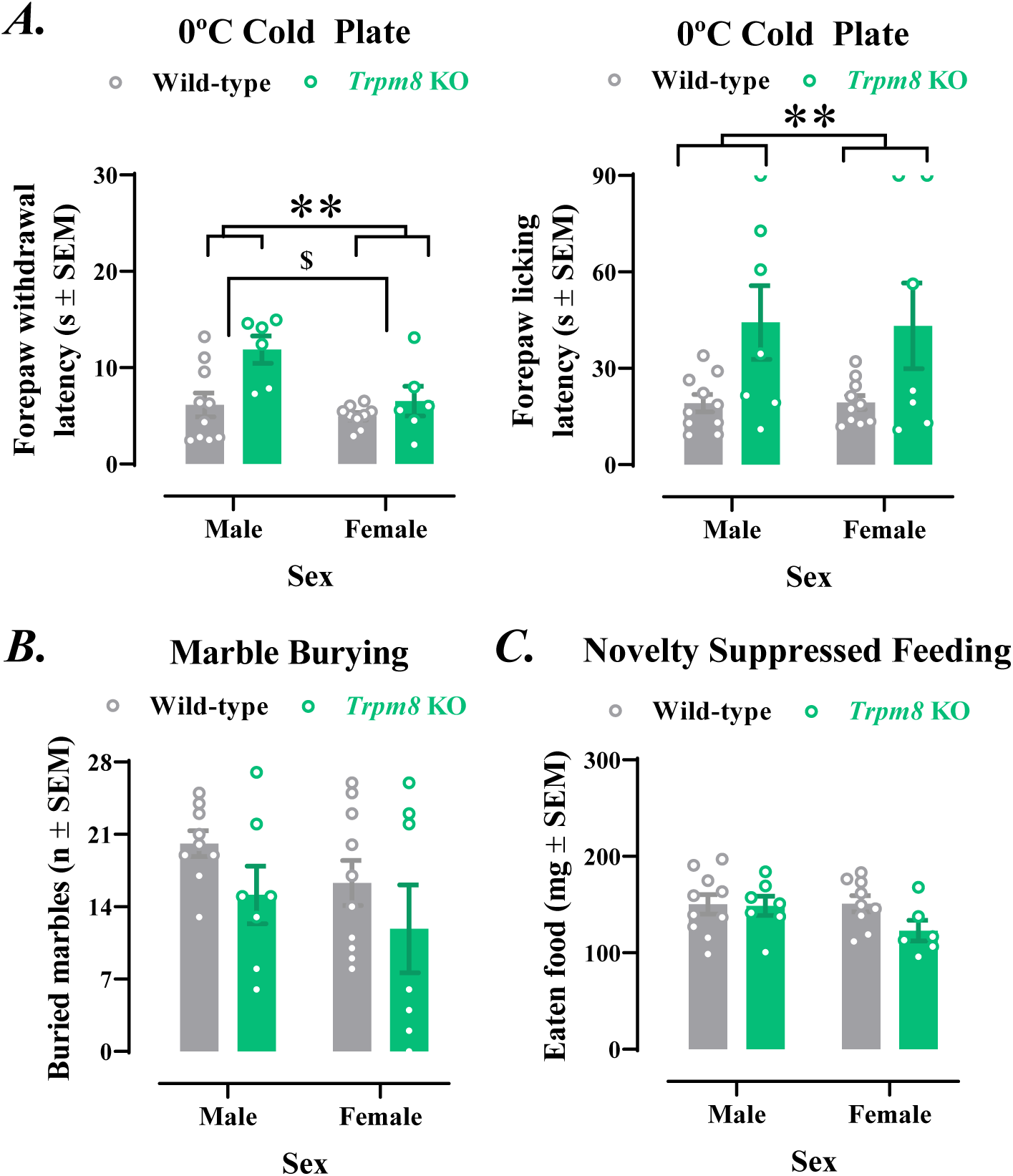
Nociceptive characterization of *Trpm8* knockout mice. A) Cold sensitivity. The Cold Plate Test at 0 °C reveals longer latencies to forepaw withdrawal (left panel) and forepaw licking (right panel) in *Trpm8* knockout mice when compared to wild-type (**P<0.01 genotype effect), suggesting a reduced cold sensitivity in knockouts. In addition, shorter forepaw withdrawal latency in females than in males (left panel, $P<0.05 sex effect) depicts enhanced cold sensitivity of females. **B) Marble Burying behaviour.** Mice buried similar number of marbles in the marble burying test regardless of sex or genotype, suggesting similar levels of defensive anxiety among wild-type and *Trpm8* knockout males and females. **C) Food intake after Novelty Suppressed Feeding** test. Wild-type and *Trpm8* knockout mice of either sex ate similar amounts of food after the Novelty Suppressed Feeding test, ruling out that increased hunger could account for the reductions in the latencies to bite the food in the novel environment. *Trpm8* KO, *Trpm8* knockout. SEM, Standard Error of the Mean. 2-way ANOVAs followed by Tukey. Dots are individual values; Error bars are SEM and bars indicate average values.

**Supplementary Fig. 2.**
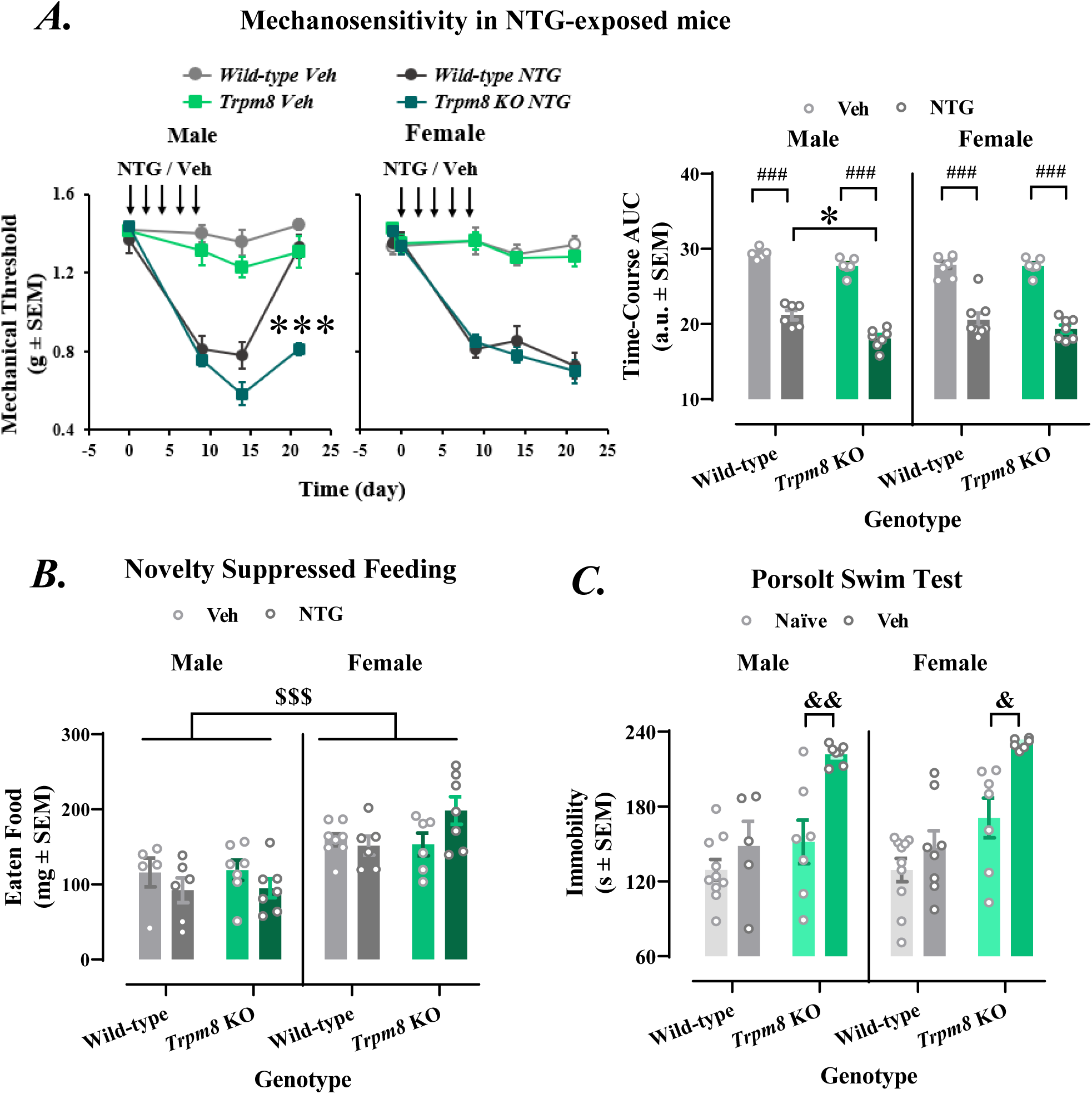
Nitroglycerin-induced mechanical hypersensitivity and effect of the experimental paradigm on *Trpm8* knockout mice. A) Nitroglycerine-induced mechanical sensitization. Wild-type and Trpm8 knockout mice develop mechanical sensitization after nitroglycerin, characterized by a decrease in the mechanical thresholds measured with von Frey filaments. This sensitization resolves 21 days later in wild-type males, but not in *Trpm8* knockout males (P<0.01 vs. wild-type males) or females of either genotype. AUCs of the time-course graphs reveal greater overall reduction of mechanical thresholds in *Trpm8* Knockout males when compared to wild-type counterparts (P<0.01) indicating stronger sensitization of the knockouts, whereas females of both genotypes behave similarly. **B) Food intake after Novelty-Suppressed Feeding.** Wild-type and *Trpm8* knockout mice ate similar amounts of food after the Novelty Suppressed Feeding test, ruling out that changes in hunger could account for the increased the latency to bite the food in knockouts exposed to nitroglycerin. Females ate more than males (P<0.001 Sex effect). **C) Depressive-like behaviour.** Control *Trpm8* knockouts receiving vehicle during the nitroglycerin model showed pronounced immobility in the **Porsolt Swim Test**, much higher than naïve *Trpm8* knockouts of previous experiments (P<0.05, males and females). This sensitivity to the experimental paradigm was absent in wild-type mice. *Trpm8* KO, *Trpm8* knockout. SEM, Standard Error of the Mean. 3-way Repeated Measures (von Frey) or 3-way (AUCs, **B**) and **C**) ANOVAs followed by Tukey post-hoc. Dots are individual values; Error bars are SEM and bars indicate average values.

**Supplementary Fig. 3.**
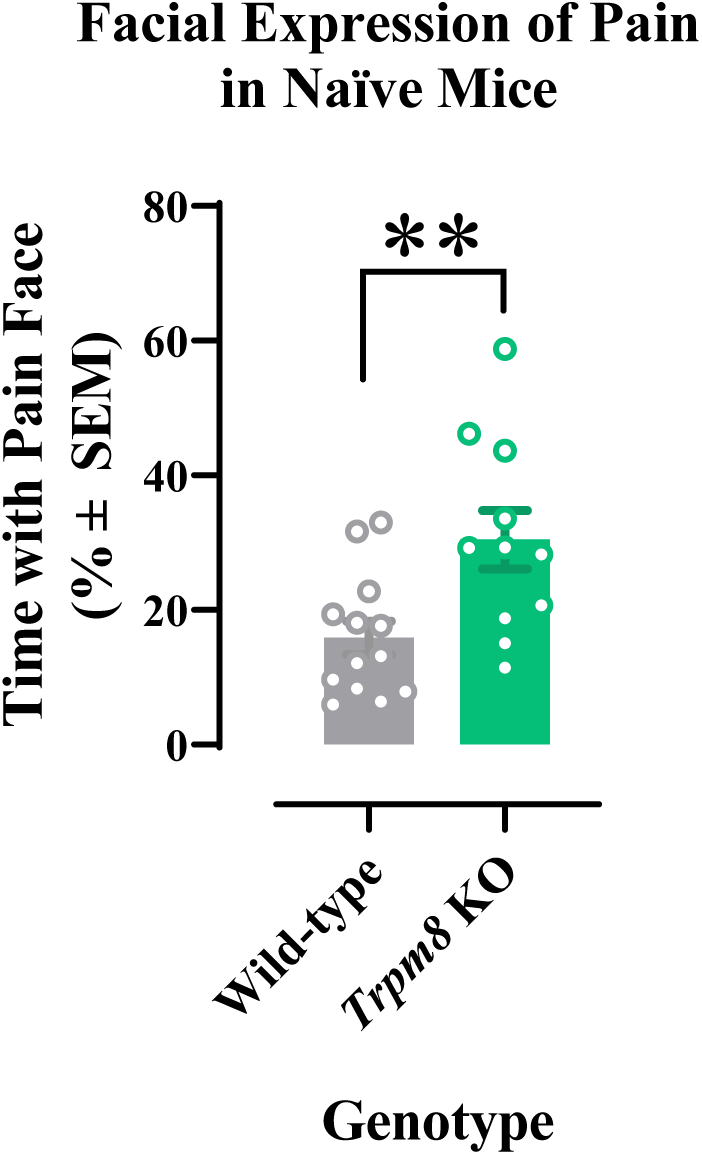
Naïve *Trpm8* knockouts displayed more facial expressions of pain than wild-type mice (P<0.01 genotype effect). SEM, Standard Error of the Mean. **P<0.01, unpaired t-test. Dots are individual values, error bars are SEM, bars are average values. *Trpm8* KO, *Trpm8* knockout.

**Supplementary Fig. 4.**
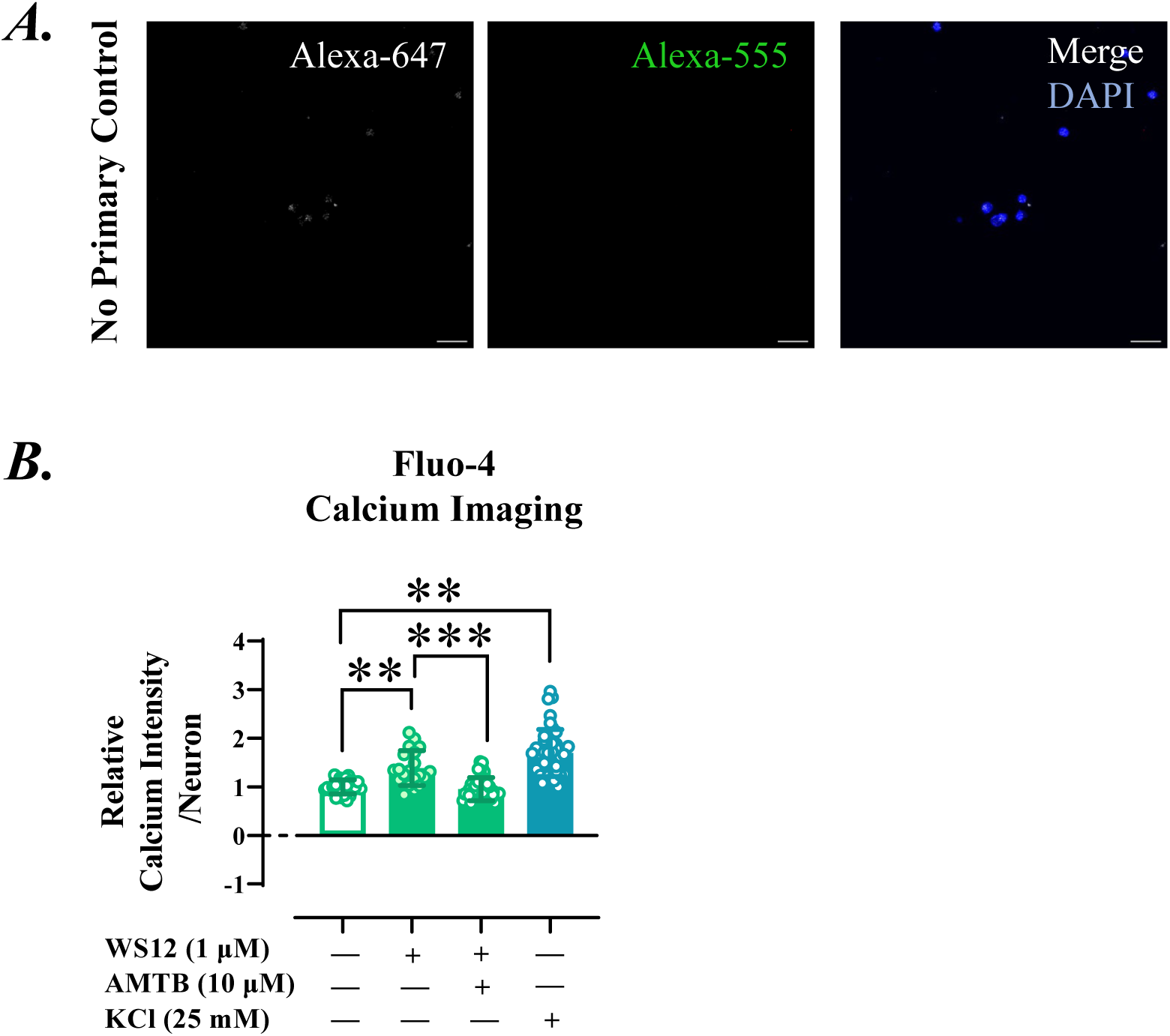
**A**) Human iPSC-derived peripheral sensory neurons lack immunoreactivity when only exposed to secondary antibodies. DAPI stains nuclei, and scale bars are of 20 µm. **B)** Fluo-4 calcium imaging conducted in human iPSC-derived sensory neurons show baseline calcium transients that are enhanced in response to 1 µM WS12, the selective TRPM8 agonist (P<0.01 vs Control). These calcium transients are sensitive to 10 µM AMTB hydrochloride, the specific TRPM8 anagonist (P<0.001 vs. WS12 without AMTB). Cells responded to 25 mM KCl, revealing their neuronal phenotype (P<0.01 vs vehicle). Dots represent individual values; error bars are Standard Deviation and bars indicate average values. ***P<0.001, One-way ANOVA followed by Tukey.

